# Primary tumor microbiomes predict distant metastasis of colorectal cancer

**DOI:** 10.1101/2025.06.02.654060

**Authors:** Bishal Parajuli, Vishal Midya, Ryan Kiddle, Nicola De Jager, Shoshannah Eggers, Daniel Spakowicz, Rebecca Hoyd, Bodour Salhia, Carlos H.F. Chan, Michelle L Churchman, Robert J Rounbehler, Song Yao, Melanie R Rutkowski, Ahmad A Tarhini, Dinesh Pal Mudaranthakam, Ashiq Masood, Therese J Bocklage, Robert W Lentz, Hassan Hatoum, Mmadili N. Ilozumba, Sheetal Hardikar, Cornelia M. Ulrich, June L. Round, Gregory Riedlinger, Craig D. Shriver, Dustin E. Bosch

## Abstract

Metastasis causes most cancer-related deaths in colorectal carcinoma (CRC), and microbiome markers may have prognostic value. We hypothesized that primary tumor microbiomes predict distant metastases. We analyzed 5-year metastasis-free survival (MFS) in a retrospective cohort of 900 ORIEN CRC tumor microbiomes (RNAseq). ORIEN findings were validated on an independent cohort using 16S rDNA sequencing and pathobiont-specific qPCR. Microbiome alpha diversity was higher in primary tumors than metastases and positively correlated with metastasis risk. Microbiome beta diversity distinguished primary vs. metastasis and predicted 5-year MFS. High primary tumor abundance of *B. fragilis* and low *F. nucleatum* were associated with short MFS. Enterobacteriaceae, including E. coli, were enriched in metastases. qPCR identified increased enterotoxigenic *B. fragilis* and *pks+ E. coli* detection in CRC metastasizers. Microbial co-occurrence analysis identified a 3-species clique that predicts metastasis (OR 1.9 [1.4-2.6]). Results suggest that primary tumor microbiomes and specific pathobionts are precision markers for metastasis risk.

## Introduction

Mortality in colorectal carcinoma (CRC) is mediated by metastasis, despite expanding multimodal treatment options^1^. Low-stage primary tumors often have excellent outcomes after surgical resection, but CRC can recur and/or metastasize several years after surgery. For example, locally advanced T-stage tumors (T2-3) without lymph node metastasis (N0) can be managed without adjuvant chemotherapy. However, follow-up is needed to identify the 10-20% of patients who will have distant metastases within 5 years.

The CRC microbiome is composed of intestinal commensal bacteria, and “pathobionts” that contribute to carcinogenesis and disease progression. Intestinal bacteria also colonize CRC distant metastases and modulate the immune microenvironment, such as tumoricidal natural killer cell activity ^2,3^. The most extensive prior research of pathobionts describes *Fusobacterium nucleatum ssp. animalis*, which adheres to host enterocytes through Fap2 and stimulates Wnt signaling through FadA interactions with E-cadherin to promote carcinogenesis and the epithelial-to-mesenchymal transition ^4,5^. In animal models, *F. nucleatum* increases the risk of metastasis and may directly contribute to lymphatic or hematogenous spread ^3,6^. *F. nucleatum* colonizes 10-20% of CRC with a predilection for right-sided location ^7^, and distant metastases ^3^. Other pathobionts have been mechanistically linked to CRC carcinogenesis and progression, such as *Escherichia coli* strains with biosynthetic genes (*pks* locus) to produce colibactin, a host DNA alkylating genotoxin ^8^. Pathobiont *pks+ E. coli* promotes carcinogenesis in the setting of colitis ^8^, and the signature mutagenic effects of colibactin are detectable in human CRC ^9,10^. Recent advances toward colibactin synthesis inhibition highlight the promise of microbiome-targeted therapies for prevention and/or treatment of CRC ^11^. Host DNA damage can also be inflicted by protein toxins, such as the type III secretion system effector UshA in *Citrobacter rodentium* and cytolethal distending toxin (CDT) in *Campylobacter jejuni* that promote carcinogenesis in mouse models ^12,13^. Some strains of the prevalent intestinal colonizer *Bacteroides fragilis* produce fragilysin or “enterotoxin” (ETBF), which promotes carcinogenesis ^14^. ETBF enhances CRC cell migration in culture, likely through a miR-139-3p-dependent pathway ^15^. ETBF also promotes “stemness” of CRC in culture through TLR4-NFAT5 ^16^.

Reflecting the importance of bacteria in CRC biology, tumor microbiomes have potential links to survival outcomes in prior research. A microbiome signature classifier with substantial contributions from *Ruminococcus spp.* was predictive of survival in several CRC cohorts ^17^. However, whether the classifier provides prognostic value beyond other dominant factors like the pathologic stage remains to be investigated. Highly powered correlational studies using the Nurses’ Health Study (NHS) and the Health Professionals Follow-up Study (HPFS) cohorts and a Chinese stage III/IV CRC cohort have suggested that *F. nucleatum* abundance predicts CRC-specific mortality (OR 1.52 [1.04-2.39]), though other studies found no correlation ^18–20^. Several known covariates with *F. nucleatum* colonization, such as right-sided anatomic location and microsatellite instability, may confound this type of single-species retrospective analysis ^21^. At least two studies have suggested that high ETBF abundance correlates with advanced stage and shorter survival ^22,23^. One mechanism by which bacteria may contribute to survival outcomes in CRC is interaction with chemotherapeutic agents. Prior research indicates that 5-fluorouracil (5-FU), a cornerstone of CRC chemotherapy since the 1980s, inhibits *F. nucleatum* growth, while tumor-resident *E. coli* can metabolize 5-FU to eliminate its anti-tumor activity ^24^.

Specific tumor-associated pathobionts have established causal roles in colorectal carcinogenesis and progression. For several organisms, rigorous prior research has identified mechanisms such as host genome modification by genotoxins and stimulation of pro-carcinogenic signaling ^25^. Knowledge gaps remain regarding the roles of *human* CRC microbiomes in disease progression, particularly metastasis risk. We hypothesized that metastasis-promoting bacteria are enriched in primary tumors of patients who eventually develop distant metastasis. We designed a retrospective cohort study using a large multi-institutional CRC population with available tumor RNAseq data. We validated the findings using 16S rDNA sequencing of an independent stage-matched validation CRC cohort.

## Materials and Methods

### Human subject cohort and study design

All human subjects research conforms to the standards of the Declaration of Helsinki. Two subject cohorts were constructed, an Oncology Research Information Exchange Network (ORIEN) discovery cohort consisting of 900 colorectal carcinoma biospecimens, and a University of Iowa Health Care (UIHC) cohort of 100 patients with colorectal carcinoma biospecimens. ORIEN is an alliance of US cancer centers established in 2014 using a standardized protocol for prospective clinical and biospecimen data collection with patients’ informed consent. Eligible patients had a tumor RNAseq dataset from at least one CRC biospecimen (primary tumor, metastasis, or both) and complete 5-year survival data (either metastasis within 5 years or at least 5 years of metastasis-free follow-up). The primary outcome was metastasis-free survival in 3 categories: 1) metastasis pre-existing at the time of biospecimen collection, 2) metastasis 0-5 years after collection, and 3) no metastasis in >5 years of follow-up. From 1315 potential candidates, 900 remained after application of the inclusion criteria. All data used in this study are de-identified with coded individual subject indicators. For the ORIEN cohort, clinicopathologic data (Table 1) were combined with microbiome relative abundance data derived from primary tumor and metastasis RNAseq with the *exotic2* pipeline for processing, contaminant removal, and normalization ^26^. Briefly, it accepts unaligned reads from tumor RNA-seq human gene expression processing (hg38 reference genome aligned with Bowtie2 ^27^) and re-aligns them to the T2T-CHM13v2.0 human reference with STAR ^28^. Unaligned reads are aligned to a custom KrakenUniq database containing finished bacterial genomes, high-quality fungal and viral genomes, as well as the T2T human reference genome and the Univec contaminants database as additional filters. Contaminants are filtered using decontam ^29^, correlating the RNA concentration to the library preparation step, and removing common contaminants in sequencing preparations ^30^.

**Table 1.**
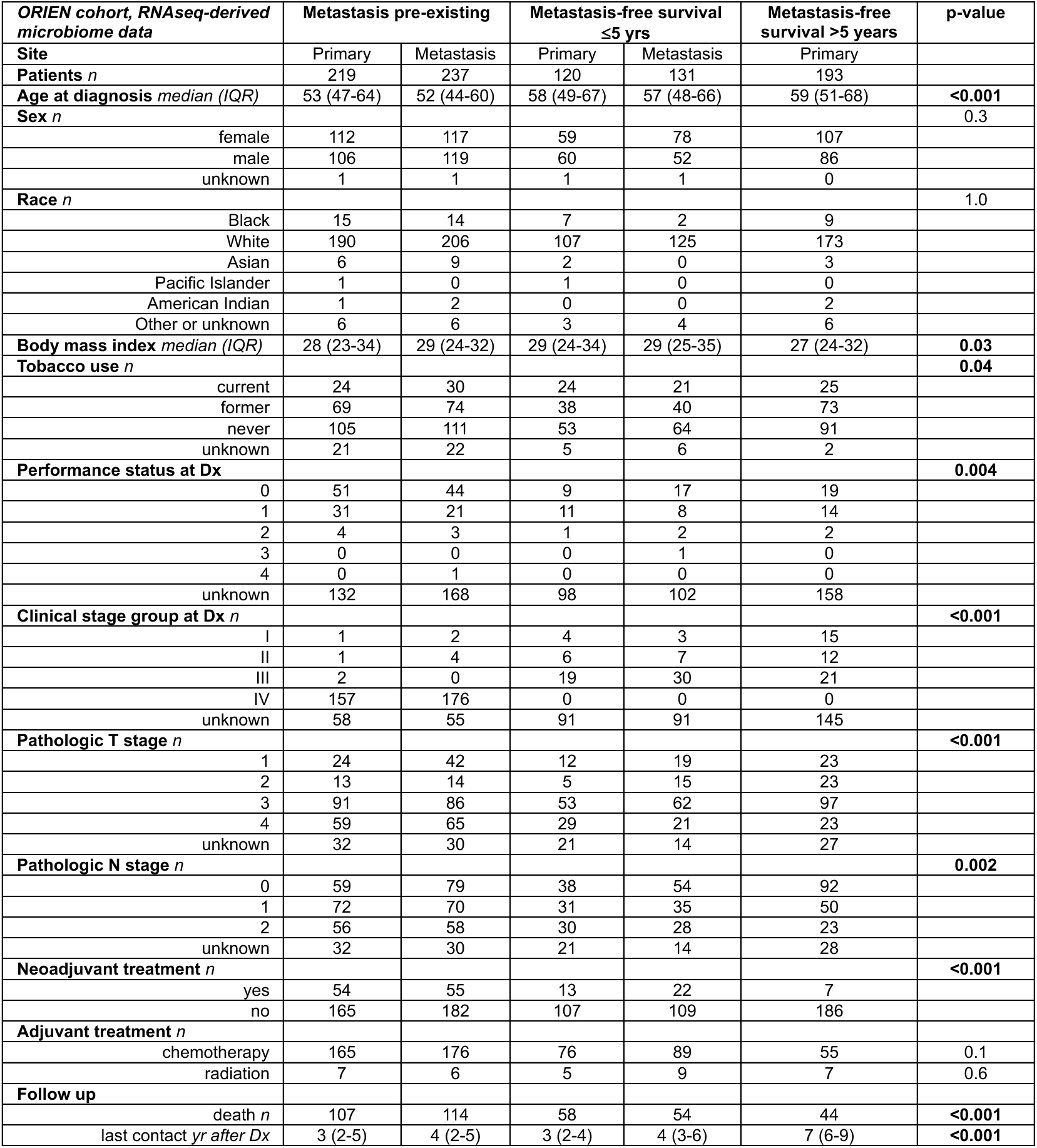
Clinicopathologic features of the ORIEN CRC cohort grouped by metastasis-free survival outcomes. All colorectal cancer patients participating in ORIEN with known metastasis-free survival (MFS) and tumor RNAseq datasets were included.

| ***ORIEN cohort, RNAseq-derived microbiome data*** | **Metastasis pre-existing** |  | **Metastasis-free survival ≤5 yrs** |  | **Metastasis-free survival >5 years** | **p-value** |
| --- | --- | --- | --- | --- | --- | --- |
| **Site** | Primary | Metastasis | Primary | Metastasis | Primary |  |
| **Patients *n*** | 219 | 237 | 120 | 131 | 193 |  |
| **Age at diagnosis median (IQR)** | 53 (47-64) | 52 (44-60) | 58 (49-67) | 57 (48-66) | 59 (51-68) | **<0.001** |
| **Sex *n*** |  |  |  |  |  | 0.3 |
| female | 112 | 117 | 59 | 78 | 107 |  |
| male | 106 | 119 | 60 | 52 | 86 |  |
| unknown | 1 | 1 | 1 | 1 | 0 |  |
| **Race *n*** |  |  |  |  |  | 1.0 |
| Black | 15 | 14 | 7 | 2 | 9 |  |
| White | 190 | 206 | 107 | 125 | 173 |  |
| Asian | 6 | 9 | 2 | 0 | 3 |  |
| Pacific Islander | 1 | 0 | 1 | 0 | 0 |  |
| American Indian | 1 | 2 | 0 | 0 | 2 |  |
| Other or unknown | 6 | 6 | 3 | 4 | 6 |  |
| **Body mass index median (IQR)** | 28 (23-34) | 29 (24-32) | 29 (24-34) | 29 (25-35) | 27 (24-32) | **0.03** |
| **Tobacco use *n*** |  |  |  |  |  | **0.04** |
| current | 24 | 30 | 24 | 21 | 25 |  |
| former | 69 | 74 | 38 | 40 | 73 |  |
| never | 105 | 111 | 53 | 64 | 91 |  |
| unknown | 21 | 22 | 5 | 6 | 2 |  |
| **Performance status at Dx** |  |  |  |  |  | **0.004** |
| 0 | 51 | 44 | 9 | 17 | 19 |  |
| 1 | 31 | 21 | 11 | 8 | 14 |  |
| 2 | 4 | 3 | 1 | 2 | 2 |  |
| 3 | 0 | 0 | 0 | 1 | 0 |  |
| 4 | 0 | 1 | 0 | 0 | 0 |  |
| unknown | 132 | 168 | 98 | 102 | 158 |  |
| **Clinical stage group at Dx *n*** |  |  |  |  |  | **<0.001** |
| I | 1 | 2 | 4 | 3 | 15 |  |
| II | 1 | 4 | 6 | 7 | 12 |  |
| III | 2 | 0 | 19 | 30 | 21 |  |
| IV | 157 | 176 | 0 | 0 | 0 |  |
| unknown | 58 | 55 | 91 | 91 | 145 |  |
| **Pathologic T stage *n*** |  |  |  |  |  | **<0.001** |
| 1 | 24 | 42 | 12 | 19 | 23 |  |
| 2 | 13 | 14 | 5 | 15 | 23 |  |
| 3 | 91 | 86 | 53 | 62 | 97 |  |
| 4 | 59 | 65 | 29 | 21 | 23 |  |
| unknown | 32 | 30 | 21 | 14 | 27 |  |
| **Pathologic N stage *n*** |  |  |  |  |  | **0.002** |
| 0 | 59 | 79 | 38 | 54 | 92 |  |
| 1 | 72 | 70 | 31 | 35 | 50 |  |
| 2 | 56 | 58 | 30 | 28 | 23 |  |
| unknown | 32 | 30 | 21 | 14 | 28 |  |
| **Neoadjuvant treatment *n*** |  |  |  |  |  | **<0.001** |
| yes | 54 | 55 | 13 | 22 | 7 |  |
| no | 165 | 182 | 107 | 109 | 186 |  |
| **Adjuvant treatment *n*** |  |  |  |  |  |  |
| chemotherapy | 165 | 176 | 76 | 89 | 55 | 0.1 |
| radiation | 7 | 6 | 5 | 9 | 7 | 0.6 |
| **Follow up** |  |  |  |  |  |  |
| death *n* | 107 | 114 | 58 | 54 | 44 | **<0.001** |
| last contact yr after Dx | 3 (2-5) | 4 (2-5) | 3 (2-4) | 4 (3-6) | 7 (6-9) | **<0.001** |

The UIHC cohort was assembled using a pathology database under a protocol approved by the University of Iowa Institutional Review Board (#202212095). Fifty patients with paired primary tumor and metastasis resections (cases) were selected. Controls with only a primary tumor resection biospecimen and at least 5-year metastasis-free survival were matched 1:1 with cases on pathologic T/N stage, age, sex, and exposure to neoadjuvant therapy. Patients with less than 5 years of follow-up and no known metastases were excluded. Eligible participants had formalin-fixed paraffin-embedded (FFPE) pathology biospecimens and slides available and complete clinicopathologic data (Table 2). Starting from 200 potential controls, 57 were excluded due to incomplete clinicopathologic data or follow-up, and 50 were selected for optimal matching to cases. Clinicopathologic data were assembled for this cohort (Table 2) using Iowa Pathology databases and the electronic health record (Epic). A board-certified gastrointestinal pathologist reviewed slides to confirm all pathologic features and select tissue at a non-neoplastic mucosal margin, the primary colon tumor, and distant metastases. Several features of this cohort were designed to complement the discovery cohort: 1) orthogonal methods for microbiome profiling using 16S rDNA sequencing and strain-specific qPCR from FFPE biospecimens, and 2) control of the known potential microbiome confounders of pathologic T/N stage and neoadjuvant therapy using matching.

**Table 2.** Clinicopathologic features of the UIHC cohort. Metastasis-free survival (MFS) groups were matched 1:1 for pathologic T and N stage and neoadjuvant therapy.

| *UIHC cohort, FFPE biospecimens* | **0-5 year MFS** | **>5 year MFS** | **test** | **p** |
| --- | --- | --- | --- | --- |
| **Patients *n*** | 50 | 50 |  | NA |
| **Age mean yr (CI)** | 54 (3) | 58 (5) | t-test | 0.1 |
| **Sex *n*** |  |  | Fisher | 0.3 |
| Female | 15 | 20 |  |  |
| Male | 35 | 30 |  |  |
| **Follow up median yr (IQR)** | 0 (0-0.5) | 5.1 (4.5-5.5) |  | NA |
| **Pathologic stage *n*** |  |  |  |  |
| *T stage* |  |  | Chi sq. | 0.3 |
| 1 | 0 | 3 |  |  |
| 2 | 4 | 9 |  |  |
| 3 | 37 | 30 |  |  |
| 4 | 9 | 8 |  |  |
| *N stage* |  |  | Chi sq. | 0.4 |
| 0 | 13 | 13 |  |  |
| 1 | 21 | 25 |  |  |
| 2 | 16 | 11 |  |  |
| X | 0 | 1 |  |  |
| *M stage* |  |  | Chi sq. | <0.001 |
| 1a | 22 | 0 |  |  |
| 1b | 4 | 0 |  |  |
| 1c | 2 | 0 |  |  |
| X | 22 | 43 |  |  |
| **Grade *n*** |  |  | Chi sq. | 0.1 |
| well differentiated | 9 | 18 |  |  |
| moderately diff. | 32 | 23 |  |  |
| poorly diff. | 9 | 9 |  |  |
| **Lymphovasc. invasion *n*** |  |  | Fisher | 0.8 |
| present | 32 | 30 |  |  |
| absent | 18 | 20 |  |  |
| **Microsatellite status *n*** |  |  |  | <0.001 |
| MSI | 0 | 11 |  |  |
| MSS | 50 | 39 |  |  |
| **Neoadjuvant therapy *n*** |  |  | Fisher | 1 |
| yes | 36 | 35 |  |  |
| no | 14 | 15 |  |  |
| **Adjuvant therapy *n*** |  |  | Fisher | <0.001 |
| chemotherapy | 49 | 40 |  |  |
| radiation therapy | 16 | 4 |  |  |

### UIHC cohort 16S metagenomic sequencing

The selected FFPE tissue was dissected from glass slides and total DNA was extracted (QIAgen FFPE Advanced UNG kit). 16S rDNA amplification, library generation, and sequencing were performed as previously described ^31^. Briefly, the V3-V4 region of the 16S rRNA gene was amplified using primers 5’-TCGTCGGCAGCGTCAGATGTGTATAAGAGACAGCCTACGGGNGGCWGCAG-3’ and 5’-GTCTCGTGGGCTCGGAGATGTGTATAAGAGACAGGACTACHVGGGTATCTAATCC-3’. Indexing and library construction were performed using a second round of PCR. Multiplexed samples were sequenced by paired-end sequencing on a MiSeq instrument (Illumina). The data were processed using QIIME 2 ^32^. Sequences were demultiplexed, and quality profiles were visualized using demux and summarize functions. The DADA2 pipeline ^33^ was used for sequence quality control and feature table generation. Phylogenetic trees were generated using the QIIME 2 phylogeny function. Alpha and beta diversity metrics with group significance statistics were calculated using the q2-diversity plugin in QIIME 2. Amplicon sequence variants were classified using a classifier trained on the Greengenes database (gg_12_8) ^34^.

In contrast with fecal metagenomics, microbiome profiling from formalin-fixed paraffin-embedded tissue (FFPE) presents unique challenges. Despite this, prior research demonstrates that bacterial communities can be accurately profiled from this matrix ^35–38^. Among the challenges are a predominance of human DNA extracted from tissue, DNA damage and cross-linking related to formalin fixation, and the potential for contaminating bacterial DNA during non-sterile specimen handling. Negative control samples included tissue-free FFPE blocks and reagents only. We applied several additional “decontamination” approaches: 1) exclusion of reads mapping to taxonomic groups that are known contaminants from FFPE processing at our institution, identified by a previous 10 subject intra-individual FFPE and fresh frozen tissue comparison, 2) statistical identification of likely contaminants with the *decontam* package in R ^29^, and 3) exclusion of taxonomic groups that are not identified in at least 1% of high-quality fecal metagenomes from the Human Microbiome Project ^39^ as measured with MetaPhlAn ^40^. After decontamination, there was a median of 14,361 high-quality reads per sample (interquartile range 7,048 – 23,975), comparable sequencing depth to previous studies of colon FFPE samples ^36^.

### Pathobiont gene quantitative PCR

Multiple qPCR primer pairs were designed for targeting *F. nucleatum nusG* and 16S rDNA, *B. fragilis* enterotoxin gene *bft1*, and *E. coli pks* locus genes *clbB* and *cnf1*. Total bacteria were quantified with primers targeting the V7 region of 16S rDNA, and human DNA using the actin gene. Primers were bioinformatically filtered for off-target effects using BLAST, NCBI databases, and a large collection of metagenome-assembled genomes (MAGs) ^41^. qPCR was performed using SYBR green and thermal melt analysis to confirm the expected amplicon. Control reactions were performed on DNA extracted from pure pathobiont cultures. Pathobiont detection in biospecimens was defined as a C_T_ value of less than 35 cycles and a thermal melt curve like pure culture controls. Detection rates were compared using chi-square testing. Relative quantities were calculated using the 2^-ΔCt^ method ^42^ with total bacteria (16S) as the referent. All qPCR primer sequences are in Supplementary Table 1.

### Microbiome data analysis and statistics

The analysis was conducted in two primary stages. The first stage includes analyses with overall diversity measures, whereas the next stage includes identifying a group of microbes, or microbial cliques, associated with the 5-year metastasis-free survival (MFS) outcome. The alpha and beta diversity metrics to be analyzed were selected before the study. Alpha diversity was quantified with Shannon’s index, Simpson’s index, and Chao1. Beta diversity was quantified with unweighted UniFrac distance, Bray-Curtis dissimilarity, and principal component analyses of both metrics. For RNAseq-derived taxonomic abundances (ORIEN), a random taxonomy-based tree was used where required; for 16S rDNA data (UIHC), a tree was constructed from the alignment of sequence clusters. Statistical testing for alpha diversity differences by site and MFS was 2-way ANOVA. Beta diversity differences were identified by the adonis2 test in R for the ORIEN dataset and the PERMANOVA test in QIIME2 for the UIHC dataset. Differential taxon abundance testing by linear discriminant analysis (LDA) was performed with LefSe using a cutoff LDA score of >2.0 ^43^. For paired primary tumor and metastasis biospecimens from the same subject, paired Wilcoxon tests were applied to alpha diversity metrics, and UniFrac distances were compared using Kruskal-Wallis tests. Gene function and bacterial trait analysis of the ORIEN dataset was performed by mapping ∼1300 detected bacteria and their relative abundances to the Protraits database ^44^. Statistical testing for trait enrichment was Mann-Whitney tests, corrected for multiple comparisons with a false discovery rate of 0.01. Gene function and pathway enrichment analysis of the UIHC 16S rDNA data was sequence-based using PICRUSt2 ^45^ and LDA. Clinicopathologic data comparisons were Chi-square or Fisher’s exact tests for categorical data and Kruskal-Wallis or Student’s t-test for numerical data. Analysis outputs were visualized using R, QIIME 2 View (view.qiime2.org), and GraphPad Prism 9 (La Jolla, CA, USA).

To discover a microbial clique associated with the outcome with a combination of an interpretable machine learning tool in a regression-based framework called the microbial co-occurrence analysis (MiCA) ^46,47^. Identifying a microbial clique consisting of multiple microbes is challenging because of the inherently large and computationally expensive combinatorial search space. MiCA utilizes a tree-based interpretable machine-learning tool to discover the most prevalent combination of microbes based on their relative abundance. Then, using a quantile-based threshold-finding algorithm creates a predictive rule associated with the outcome. Based on the relative abundances of the most frequently co-occurring microbes, this predictive rule intuitively creates a mathematical equivalence of a microbial clique. Lastly, we use a covariate-adjusted regression to estimate the effect of this microbial clique. Since machine-learning tools naturally tend to overfit, we trained the tree-based algorithm in MiCA using 65% of the data (n = 769). Then the remaining 35% of the data (n=416) was used to validate the identified microbial clique. This two-stage procedure ensured the robustness of the identified microbial clique. The regression part of the MiCA algorithm incorporates multiple covariates, including age at diagnosis, reported sex, race, neoadjuvant therapy (yes/no), clinical stage group at biospecimen collection, and pathologic stage group. The dominant clique identified includes *Escherichia coli*, *Bacteroides fragilis*, and a *Pseudomonas sp.* ASV. The Kraken database and algorithm (the highest stringency parameter for taxon assignment was used) assigned this ASV as *Pseudomonas yamanorum*, which is neither a common gut commensal nor a common environmental contaminant. BLAST of several sequences from the taxonomic unit showed 100% identity to *P. yamanorum* genomes and >90% identity to at least five other *Pseudomonas spp.* Since we believe this could be an over-call of the species-level classification by Kraken, we refer to it as a *Pseudomonas sp.* in the results.

## Results

### Descriptive statistics of independent discovery and validation cohorts

Uniformly tabulated clinicopathologic data from the ORIEN cohort showed younger age in the pre-existing metastasis group, modest differences in BMI (lowest in >5-year MFS), slightly more current smokers in the 0-5 yr MFS group, and differences in known vs. unknown performance status (Table 1). Since the primary tumor stage is a dominant predictor of MFS ^48^, the higher pathologic and clinical stage parameters and death rates among the groups with metastasis were expected. Neoadjuvant therapy is more commonly indicated and given in the advanced stage and short MFS groups. The length of follow-up difference was expected due to study design, because only the >5-year MFS group was required to have at least 5 years of follow-up.

Clinicopathologic data from the UIHC cohort showed more frequent microsatellite instability in the >5 yr MFS group (Table 2). The M stage was specified by study design. Adjuvant chemotherapy was more frequently given to metastasizers, as expected. Since adjuvant therapy was given *after* biospecimen collection in all cases, it does not affect the primary tumor microbiome.

### CRC tumor microbiome diversity predicts metastasis-free survival

We first asked whether bacterial communities colonizing primary tumors are related to the risk of metastasis. To address this question, we compared tumor microbiomes from patients with metastasis preceding the biospecimen collection (pre-existing metastasis), those that metastasized within 5 years, and those with more than 5 years of MFS. Primary tumor microbiome alpha diversity was slightly lower in patients with long-term MFS, having the same trend and borderline statistical significance with multiple metrics in the discovery and validation cohorts (Figure 1A-C, 2-way ANOVA p 0.02-0.08). All analyses were stratified by site (metastasis vs. primary tumor) and metastases consistently showed lower alpha diversity. Tumor microbiome beta diversity analysis also showed significant differences between the MFS groups, as well as sites (Figure 1D-E). All the above global diversity trends were found in both cohorts. The ORIEN discovery cohort has several clinicopathologic differences among the MFS groups that could confound the microbiome associations with disease progression. Since the cohort is large enough to power multivariate modeling, we asked whether the MFS-microbiome link is independent of several potential confounders using a permutational clustering approach implemented in adonis2 (Table 3). A statistically significant correlation between microbiome beta diversity and MFS persisted after adjustment for potential confounders.

**Figure 1.**
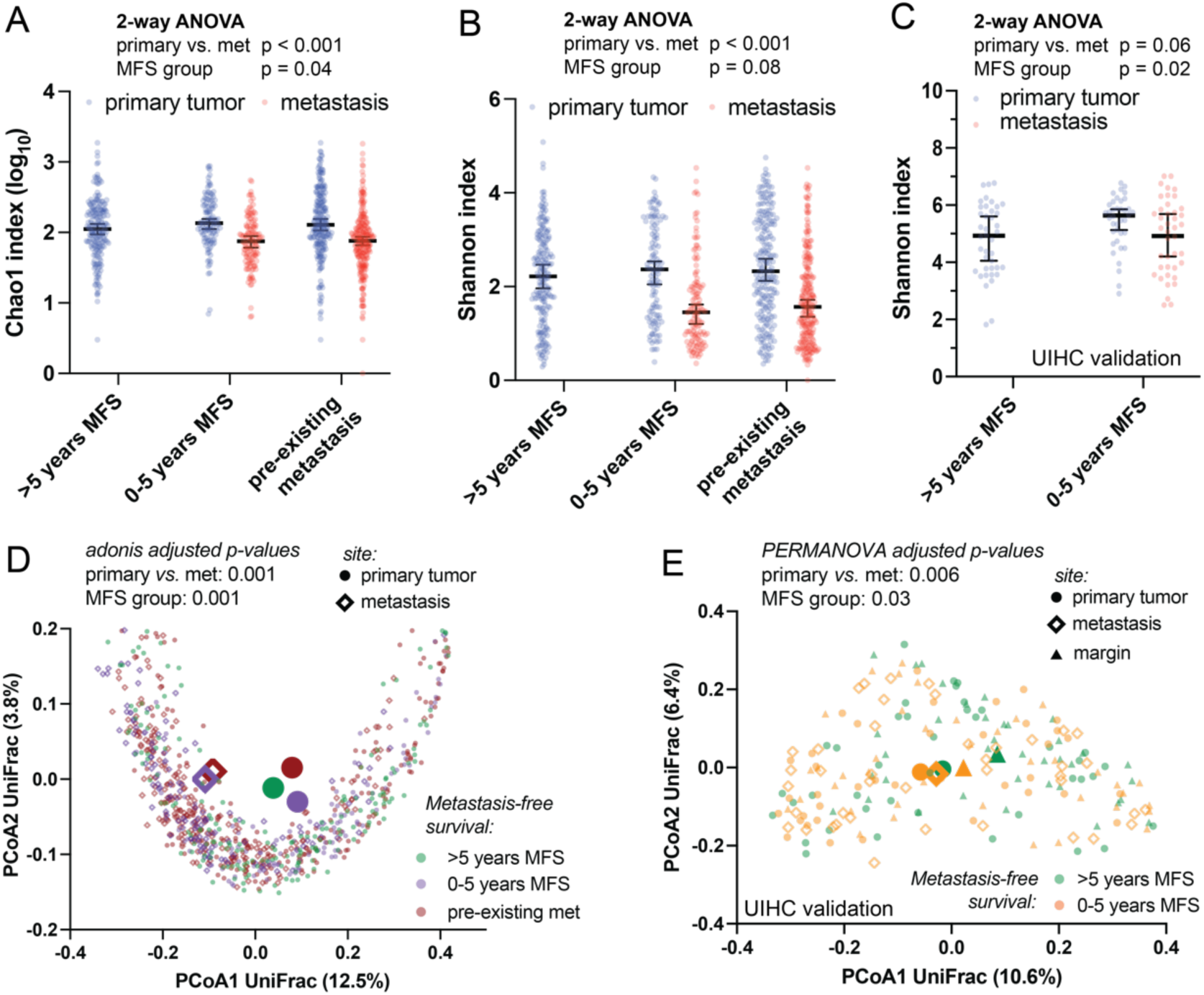
CRC tumor microbiomes predict metastasis-free survival. (A, B) In the ORIEN cohort, alpha diversity by several metrics is significantly lower in distant metastases, and among primary tumors, modestly lower in patients with long term metastasis-free survival (MFS). (C) Similarly, low alpha diversity in primary tumors correlated with shorter MFS in the UIHC cohort. Metastases had lower Shannon indices that did not reach statistical significance. Microbiome beta diversity in both the ORIEN (D) and UIHC (E) cohorts is also distinguishable by site (metastasis vs. primary tumor) and MFS group. Larger symbols represent calculated centroids. The findings indicate that tumor microbiome diversity corresponds to MFS, and therefore may have prognostic value.

**Table 3.**
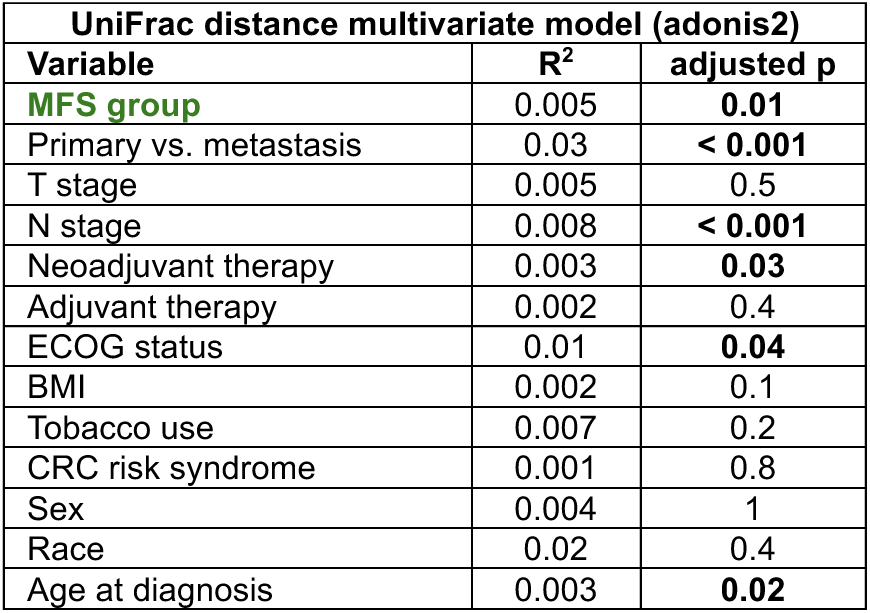
Tumor microbiomes still predict MFS after adjustment for potential confounders. Adonis statistical testing of beta diversity relationships to MFS were performed with adjustment for potential confounders that differ by MFS study group in the ORIEN cohort (Table 1). The association between beta diversity and MFS was not explained by co-variance with site, age at diagnosis, regional lymph node status, or neoadjuvant treatment, although many factors tested were significantly related to beta diversity.

To identify specific taxa that distinguish metastasis risk, some of which may be causally related to disease progression and thus therapeutic targets, we applied LefSe ^43^ linear discriminant analysis (Figure 2). Among the most discriminating features in the ORIEN dataset were at least six known pathobiont-containing taxa that contribute to carcinogenesis through known mechanisms (Figure 2A, blue text). Surprisingly, *F. nucleatum* was highest in primary tumors from patients with long-term metastasis-free survival. This may be explained by an inverse correlation between *F. nucleatum* abundance and stage group, *i.e., F. nucleatum* is most abundant in tumors with a shallow depth of invasion and a lack of lymph node metastases (Kruskal-Wallis p = 0.003). Analysis of the stage-matched UIHC validation cohort revealed different discriminating taxa, although some overlap with the discovery cohort was seen (Figure 2B, red text). Thus, beta diversity differences according to MFS are strongly consistent, but the specific discriminating taxa differ between the two cohorts, which could be due to covariance with their distinct clinicopathologic features (Tables 1-2).

**Figure 2.**
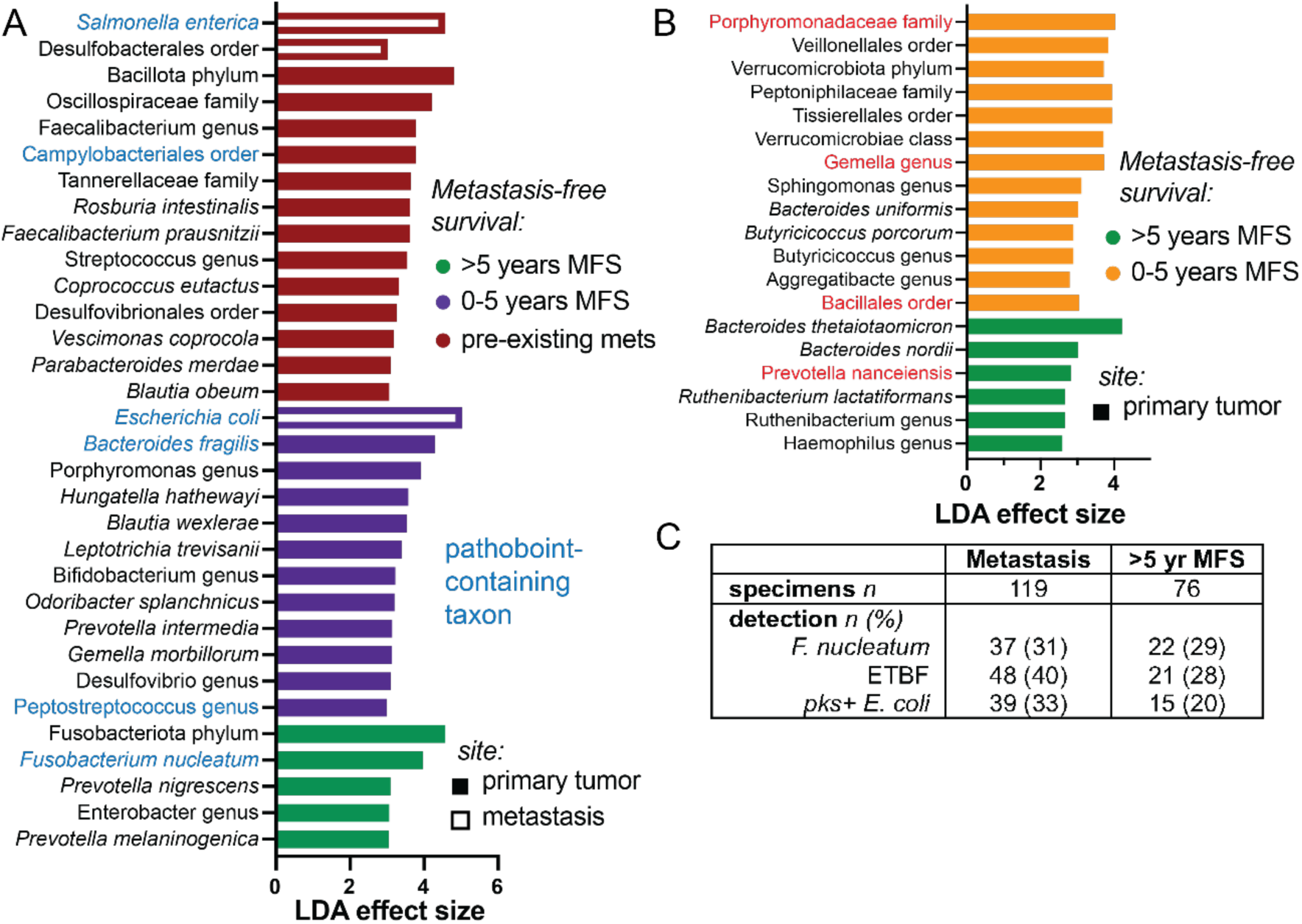
Pathobiont taxa predict CRC metastasis-free survival. (A) Linear discriminant analysis of the ORIEN cohort identified several distinguishing taxonomic groups that contain known CRC-related pathobionts (blue). The Enterobacteriaceae *S. enterica* and *E. coli* are enriched in distant metastases. *B. fragilis* and *Peptostreptococcus spp.* in primary tumors predicted metastasis within 5 years, while *F. nucleatum* associated with long-term MFS.(B) Discriminating taxa in the stage-matched UIHC cohort were not the same, but overlapping taxonomic groups are highlighted in red. (C) Tissue RNAseq- and 16S-derived microbiome data does not distinguish pathobiont strains, so qPCR for *F. nucleatum*, *bft1-3* from enterotoxigenic *B. fragilis* (ETBF), and *cnf2* and *clbB* genes from the *pks* locus in colibactin-producing *E. coli* were performed on tumors and non-neoplastic margins. Detection rates were significantly different (Chi-squared p < 0.01) with ETBF and *pks+ E. coli* more frequently detected in metastasizer biospecimens. *F. nucleatum* detection rates did not differ in the UIHC cohort by MFS group.

The 16S rDNA sequencing cannot resolve specific pathobiont strains, and the RNAseq data were of insufficient depth to confidently discriminate the closely related genomes. For example, 16S rDNA sequences do not distinguish enterotoxigenic from nonenterotoxigenic *B. fragilis*. We selected three of the most studied CRC-promoting pathobionts for strain-specific detection and quantitation by PCR: *F. nucleatum*, enterotoxigenic *B. fragilis* (ETBF), and *pks+ E. coli*. qPCR performed on all non-metastasis biospecimens from the UIHC cohort (primers listed in Table S1) showed more frequent detection of both ETBF and *pks+ E. coli* in metastasizers than patients with long-term MFS (Figure 2C, 33-40% vs. 20-28%, Chi squared p < 0.01). Detection of *F. nucleatum* was not associated with MFS in the stage-matched UIHC cohort.

### Microbial co-occurrence predicts metastasis-free survival

We identified a microbial clique of the simultaneous presence of *Escherichia coli* and the absence of a *Pseudomonas sp.* & *Bacteroides fragilis* in the training data (randomly selected 65% of the samples). Almost half (47.6%, n = 565) of the total study sample had this microbial clique. This microbial clique was strongly associated with increased odds of metastasis (OR [95% CI]: 1.91 [1.40,2.59], p-value < 0.0001) across the whole sample (Figure 3). It was also strongly associated with increased odds of metastasis (OR [95% CI]:1.88 [1.11,3.20]) in the test data (35% of the sample).

**Figure 3.**
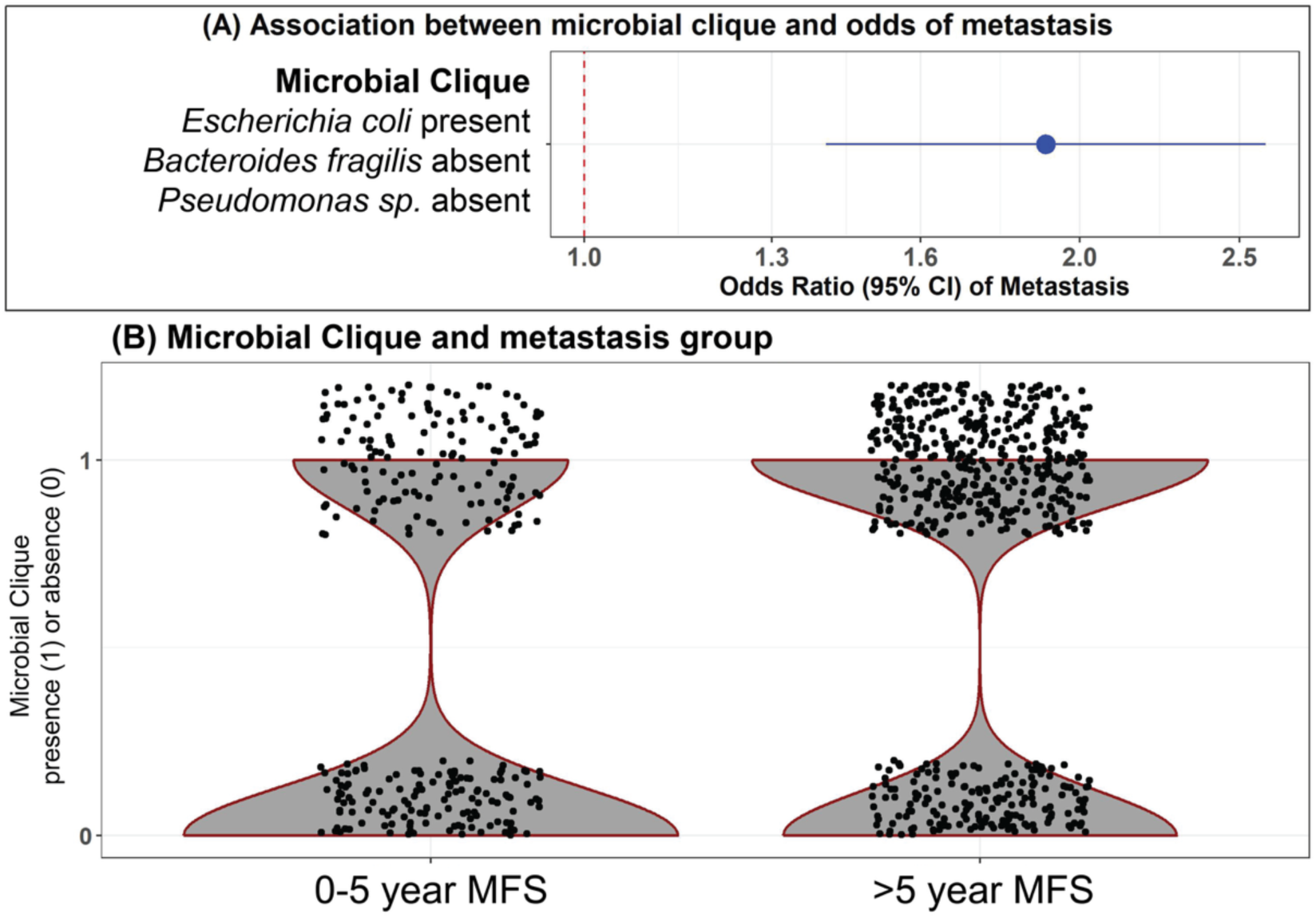
A microbial clique identified with the MiCA algorithm can predict metastasis. (A) A forest plots shows the association between the identified microbial clique and the odds of metastasis. (B) A violin plot shows the distribution of the outcome among patients with and without the clique.

### Primary tumor microbiome capacities for carbohydrate metabolism and quinone biosynthesis reflect metastasis risk

Although specific MFS-associated taxa, other than pathobionts, did not show a high degree of concordance between the two cohorts, we reasoned that the functional capacities of the microbiome community may correspond to tumor microenvironment differences underlying metastasis risk. These functions may also highlight targets for future mechanistic studies and microbiome-targeted interventions. Using predicted gene function in the UIHC cohort, we examined correlations with MFS (Figure 4A). The dominant trends were the enrichment of certain carbohydrate catabolism pathways and quinone biosynthesis pathways in tumors from patients with long-term MFS. Several enriched pathways are involved in the biosynthesis of menaquinones, which are components of the cell membrane that participate in electron transport. Menaquinones are prevalent, reversible redox components in anaerobes and most Gram-positive aerobes ^49^. For the ORIEN cohort, we leveraged a large database of phenotype inferences, ProTraits ^44^, and mapped ∼1300 of the detected taxa in our biospecimens to derive predicted trait abundances. A comparison of the 0-5 yr and >5 yr MFS group microbiomes revealed a few traits significantly enriched in the long-term MFS group (Figure 4B). Among these were several carbohydrate utilization traits and strict anaerobic preference, which correspond well to the metabolism and anaerobe-enriched menaquinone biosynthesis in the UIHC cohort.

**Figure 4.**
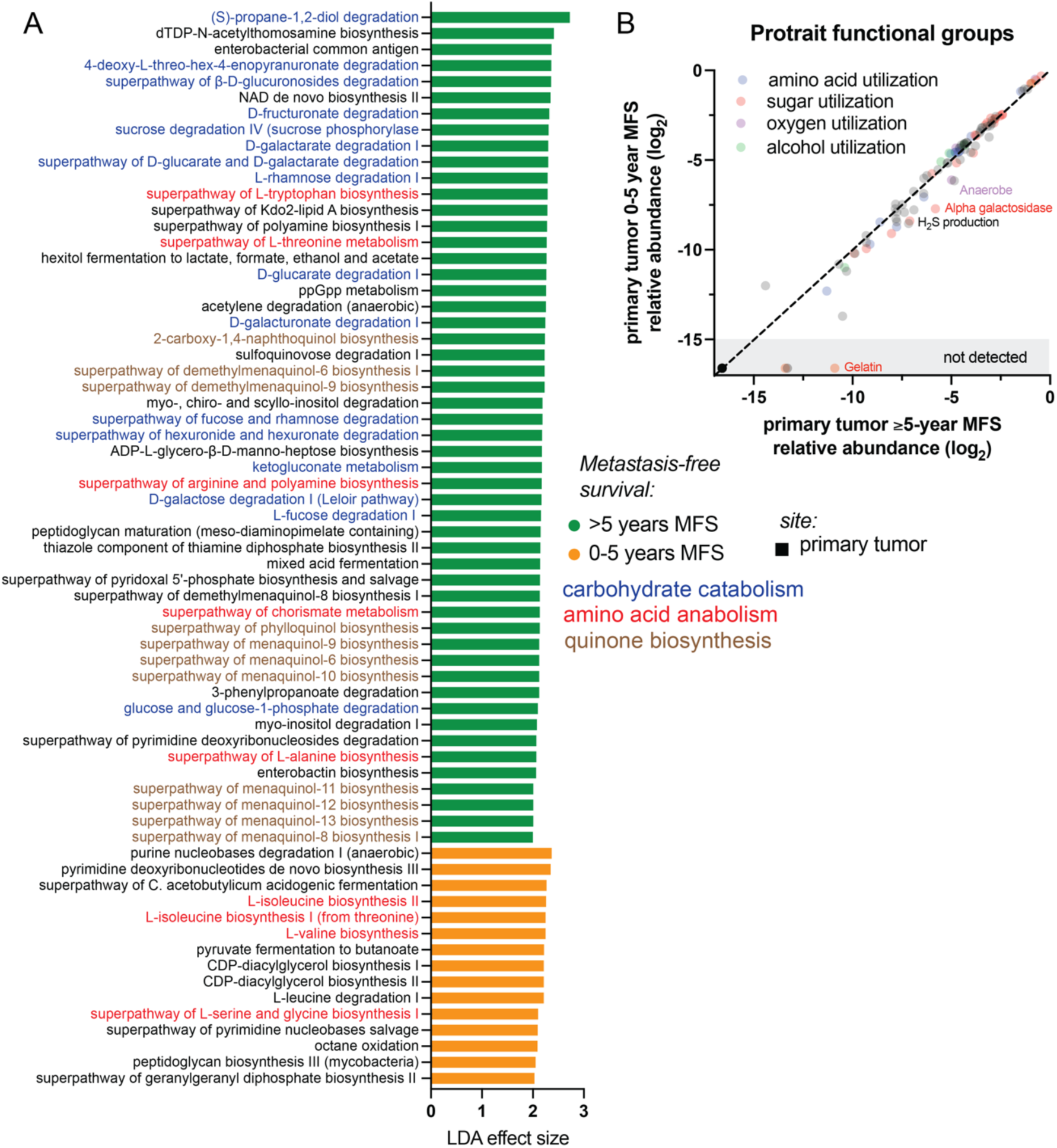
Predicted primary tumor microbiome functional capacities are related to metastasis risk. (A) For the UIHC cohort, linear discriminant analysis was performed on 16S rDNA sequence-based pathway abundance prediction from Picrust2. Several patterns emerged as enriched in the > 5-year MFS group: carbohydrate catabolism pathways (blue), quinone biosynthesis (brown), and some amino acid biosynthesis pathways (red). The most enriched pathways in metastasizer primary tumors were nucleic acid metabolism pathways. (B) For the marker gene-derived microbiome data in the ORIEN cohort, the identified species were mapped to the Protraits bacterial traits database. Among the ∼1300 species with phenotypic data, few specific associations were found related to MFS: strict anaerobes, alpha galactosidase, and hydrogen sulfide producers were modestly enriched in the >5-year MFS group.

### Metastasis microbiomes derive from the primary tumor and have reduced alpha diversity

CRC metastases are colonized by intestinal bacteria including pathobionts ^3^. We assessed correlations between the microbiomes of primary tumors and metastases through paired biospecimens from individual patients. Consistent with our observation that metastasis microbiomes in general have lower alpha diversities (Figure 1A-C), alpha diversity was lower in metastases compared to the paired primary tumor from the same subject in both the ORIEN and UIHC cohorts (Figure 5A-C). To assess beta diversity relationships between primary/metastasis pairs, we quantified UniFrac distances. In the ORIEN cohort, paired primary tumor and metastasis microbiomes within one subject were more closely related than unpaired comparisons (Figure 5D). This pattern did not hold in the UIHC validation cohort, though statistical power was limited by the number of paired biospecimens (Figure 5E). Overall, the findings suggest that similar bacterial populations colonize both primary tumors and metastases within an individual. However, a restricted set of taxa (lower alpha diversity) can colonize and persist at distant sites.

**Figure 5.**
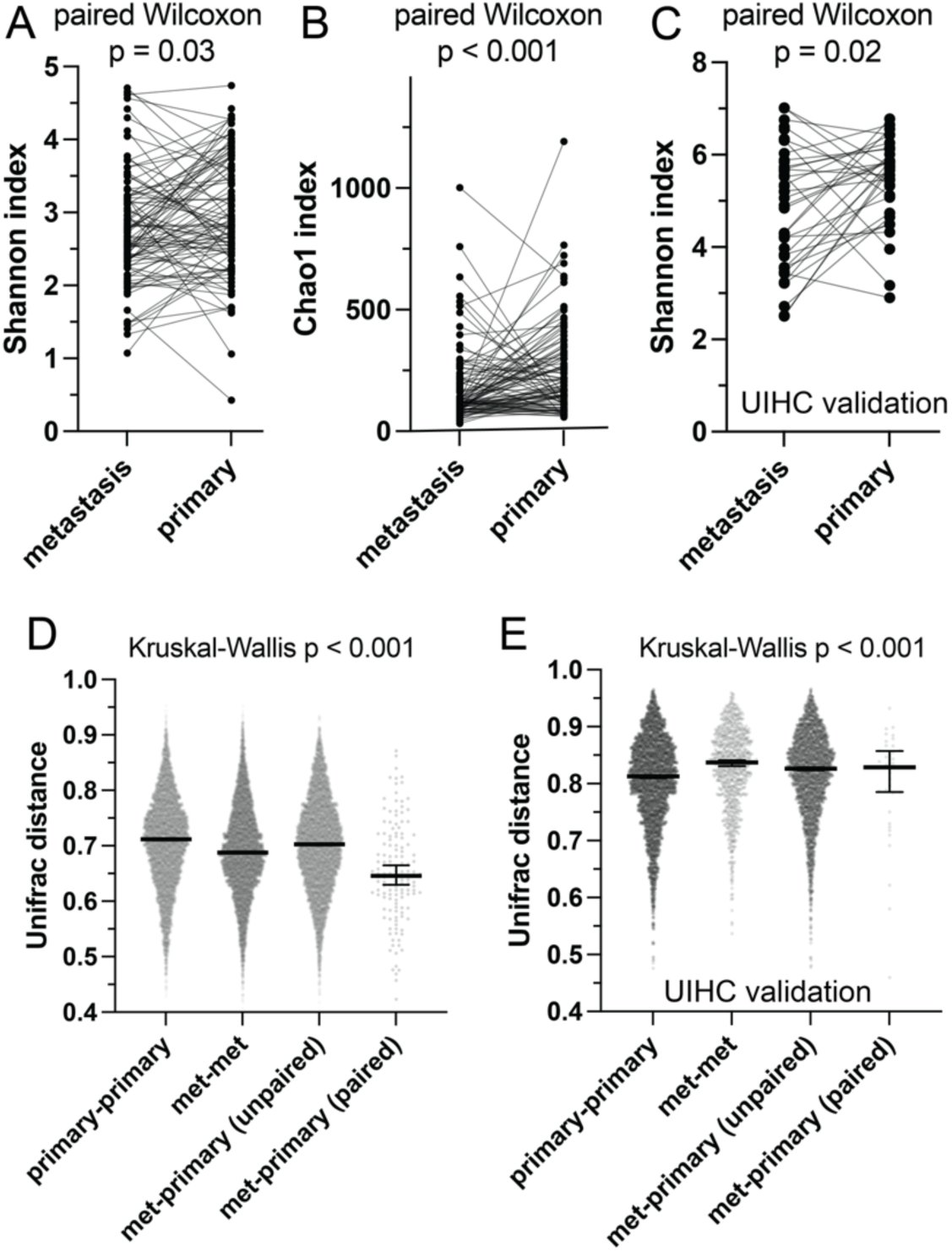
Metastasis microbiomes are closely related to the primary tumor microbiome but with reduced alpha diversity. Paired primary tumor and metastasis biospecimens with microbiome data were available for 103 patients in the ORIEN cohort. (A B) A paired analysis demonstrated lower alpha diversity in distant metastases, a finding validated in the stage-matched UIHC cohort (C). (D) ORIEN cohort paired metastasis and primary tumor microbiomes were closely related by beta diversity distance metrics in contrast with unpaired biospecimens, compatible with similar bacterial communities from the primary tumors colonizing distant sites. Inter-subject primary tumors were the most distant. (E) However, this pattern was not seen in the UIHC cohort, where primary tumor microbiomes were most similar.

We then examined differentially abundant taxa in the paired primary tumor and metastasis microbiomes. Several taxonomic groups were significantly depleted in metastases compared to primary tumors (Figure 6A-B). The facultative anaerobe and pathobiont-containing Bacteroidaceae family and Fusobacterium genus, as well as the Bifidobacterium genus, were significantly less abundant in metastases from both cohorts. Enterobacteriaceae, including *E. coli,* had similar abundances at both sites. The species *Fusobacterium periodonticum* and a Bacillus genus ASV were significantly enriched in the metastases of the UIHC cohort. The findings suggest that some major microbiome families, predominantly anaerobes, are less fit for survival in distant metastasis microenvironments. ProTraits functional group analysis (Figure 6C) identified strict anaerobicity and several carbohydrate and amino acid utilization traits as depleted in metastases. Aerobes, facultative anaerobes, and alcohol utilization traits had modest advantages in metastases. Analysis with PICRUSt2 in the UIHC cohort identified few pathways enriched in metastases (none in the primary tumors), with themes of nucleic acid metabolism, quinone biosynthesis, and peptidoglycan biosynthesis (Figure 6D). The findings suggest that nutrient sources (carbohydrates and amino acids) and oxygen levels in distant metastasis microenvironments negatively select some of the taxa that colonize primary tumors.

**Figure 6.**
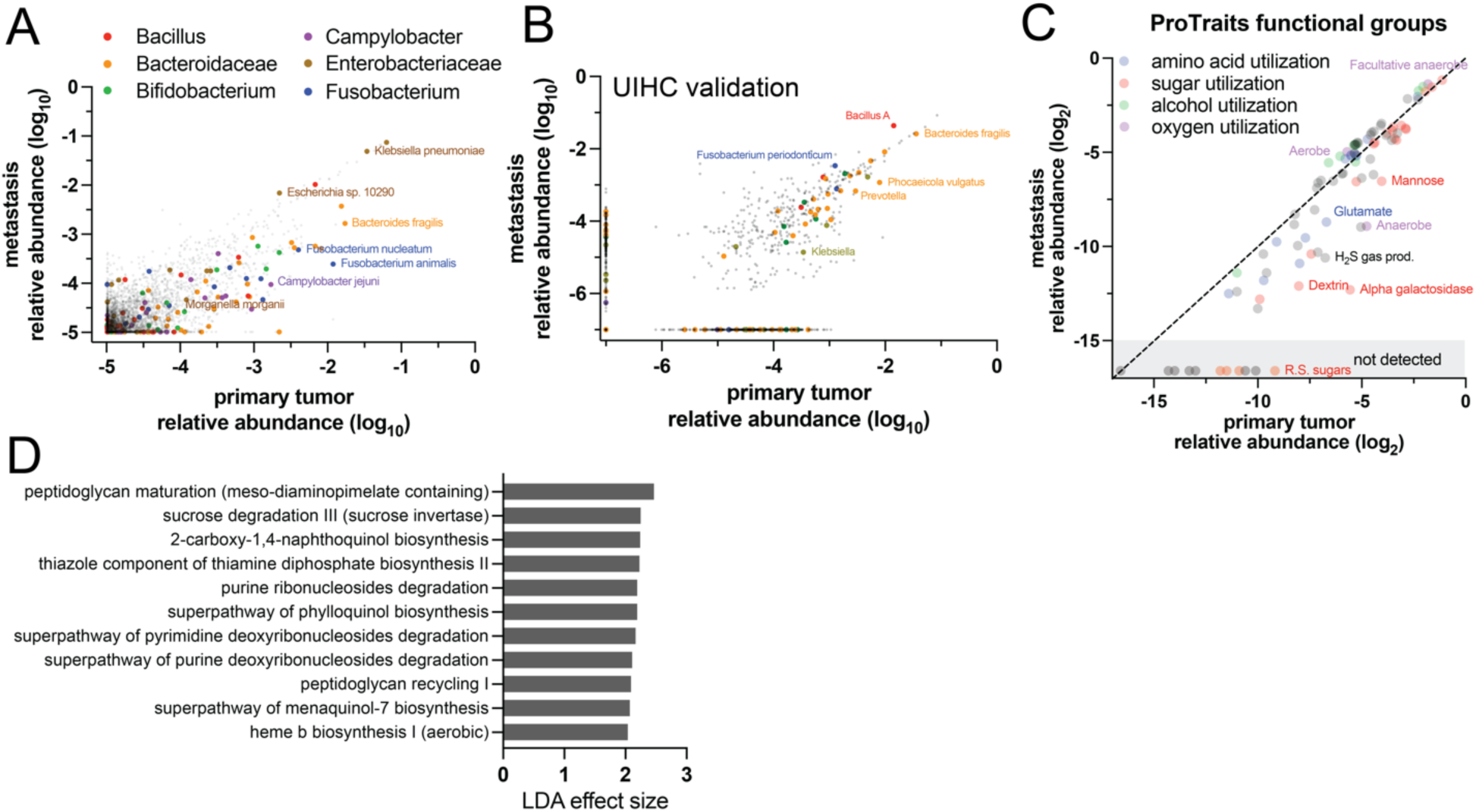
Colonization of metastases by tumor-associated bacteria is related to oxygen tolerance and metabolic capacities. (A) In the ORIEN cohort, relative abundance comparison to primary tumors showed metastasis depletion of *Fusobacterium spp.,* Bacteroidaceae, and *Bifidobacterium spp.*, among others, while Enterobacteriaceae and *Bacillus spp.* were unchanged or enriched. (B) A similar analysis in the UIHC cohort identified similar patterns, except that *Fusobacterium* was not significantly depleted. (C) The Protrait database was used for enrichment analysis of bacterial functions based on taxonomy and relative abundance in the ORIEN cohort. Anaerobic bacteria were depleted in metastases, as well as H_2_S-producers and bacteria with the capacity to utilize certain sugars. The findings may reflect key selective pressures in the metastasis microenvironment, which tends to be oxygenated and likely lacks the diet-derived carbon sources available to bacteria in the gut. (D) Picrust2 predicted pathway analysis of the UIHC cohort identified modest enrichment of nucleic acid metabolism and peptidoglycan-related pathways in metastasis microbiomes. No pathways were significantly enriched in primary tumors.

## Discussion

The main finding of this study is that primary CRC tumor microbiome diversity, gene function capacities, and microbial cliques are predictive of metastasis-free survival. While most prior detailed mechanistic investigations have focused on the microbiome’s contributions to initiating carcinogenesis, we highlight that human tumor-resident bacteria also associate with disease progression and can help predict the critical clinical outcome of distant metastasis. Several pathobiont taxa, including *pks+ E. coli* and ETBF, are known to promote carcinogenesis and, in limited studies, have been shown to promote epithelial-mesenchymal transition^15,16,50^, are among the bacteria enriched in primary tumors that progress to distant metastasis. In a recent global study of CRC mutation signatures, colibactin-associated mutations showed regional variability and enrichment in early-onset colorectal carcinoma ^10^. *B. fragilis* can promote chemoresistance through host Notch1 signaling, independent of enterotoxin ^51^. The capacities of these pathobionts to mutagenize host cells, stimulate pro-proliferation and -invasion signaling pathways, promote chemoresistance, and manipulate tumor immune cells likely contribute to metastasis risk. Thus, some mechanistic insight for bacterial promotion of CRC metastasis already exists. However, further study is needed to fully understand these mechanisms and identify causal relationships between other bacteria and metastasis. Ongoing efforts to inhibit *B. fragilis* enterotoxin and colibactin biosynthetic enzymes highlight a promising opportunity to target pathobionts in CRC ^11,52,53^.

Our findings indicate that primary tumor microbiomes, pathobiont qPCR markers and specific microbial cliques have prognostic value for metastasis risk, independent of the pathologic stage at resection ^54^. The pathobiont-containing species *E. coli* and *B. fragilis* are central to the predictive microbiome features in three parts of the analysis: 1) linear discriminant analysis by MFS outcomes, 2) pathobiont strain detection, and 3) their inclusion in a highly MFS-predictive microbial clique. These features may be useful prognostic markers for determining post-resection surveillance and decisions to treat stage II CRC with adjuvant therapy. While tumor microbiome metagenomics is not part of current clinical practice, detection of specific metastasis risk-related pathobionts such as *pks+ E. coli* and ETBF by qPCR and/or *in situ* hybridization in CRC pathology specimens is imminently feasible as a clinical test ^55^. Further study is needed to develop these tests and determine their potential clinical utility as precision oncology markers.

The second significant finding of this study is that CRC metastasis microbiomes have a composition that is related to the primary tumor but with lower alpha diversity and depletion of specific bacterial traits. Microbiome capacities for carbohydrate utilization and anaerobic electron transport in tumor microbiomes are associated with long-term metastasis-free survival. One potential explanation for lower anaerobe abundances in metastasizing tumors could be greater oxygen content (less hypoxia) within the tumor microenvironment that selects against anaerobes. Bacteria in beneficial tumor microbiomes (patients with long-term MFS) have a high capacity for specific carbohydrate utilization, indicating the potential for dietary interventions to influence metastasis risk. The tissue microenvironments in the liver and lungs likely exert selective pressures on the CRC-associated microbiome due to factors such as high oxygen tension, less diverse food-derived carbon sources, and distinct immune cell populations. Prior research indicates that bacteria within metastases can influence tumor growth and response to therapy ^3,6,56^. While the prior research in this area has focused almost exclusively on *F. nucleatum*, we find that Fusobacteriaceae and several other pathobiont-containing taxonomic groups are depleted in metastases compared to the primary tumor from the same patient. Understanding the drivers of microbiome composition in distant metastases may provide insight into bacterial roles in disease progression and highlight potential interventions such as dietary changes and selective antibiotics directed at tumor microbiomes.

An outstanding goal in the field of tumor microbiome studies is intervention with microbiome-targeted therapies. Our study suggests several avenues for further investigation toward this long-term goal. First, antimicrobials or biosynthetic gene inhibitors directed at some of the known CRC carcinogenic pathobionts (colibactin-producing *E. coli* and ETBF) ^11,53^ are likely to also favorably impact disease progression. The differential capacity for carbohydrate and amino acid metabolism in primary tumors that metastasize suggests that it may be possible to manipulate the tumor microbiome with dietary (prebiotic) interventions. Dietary patterns such as the sulfur microbial diet are known to modify both the fecal microbiome and the risk of CRC ^57^, but little is currently known regarding dietary effects on tumor-resident bacteria and disease progression. Finally, while this study emphasizes pathobionts, tumor bacteria associated with long-term MFS, such as *B. thetaiotaomicron* and *Prevotella spp.* (Figure 2B) are candidate probiotics for metastasis prevention.

### Limitations

All tumor microbiome studies work from low bacterial biomass specimens, which, together with non-sterile handling of pathology specimens, increases the risks of contamination by exogenous bacterial DNA compared to fecal microbiome studies. These risks are mediated by the inclusion of controls and bioinformatic decontamination approaches (see methods). Although some degree of contamination undoubtedly remains, it is expected to be biased toward the null hypothesis (high variability, no difference between groups) because the study design compares biospecimens handled in the same way and is not expected to have systematically different contamination risks. Perhaps the greatest strength of this study is the use of two independent cohorts with different methodologies (RNAseq and 16S rDNA metagenomics), yielding highly consistent findings. As a retrospective biospecimen study, the results are primarily correlative and, in isolation, do not allow conclusions about the causal roles of the microbiome in metastasis. However, we highlight several organisms with known mechanistic causal roles in CRC carcinogenesis and progression that also emerged in our study, supporting the conclusion that some of the correlations do have an underlying causal relationship.

## Data availability

Data from the Oncology Research Information Exchange Network (ORIEN) is accessible through project requests. All 16S rDNA sequencing data will be made available through NCBI SRA, BioProject PRJNA1244713 upon publication.

## Conflict of interest statement

RWL acknowledges clinical research funding to institution from ALX Oncology, Merck, Lilly, Guardant, EDDC, Boehringer Ingelheim, and AstraZeneca, consulting/advisory role (honoraria to institution) for Agenus, Boehringer Ingelheim, Merck, Natera, and consulting/advisory role (unpaid) for Kahr and EDDC. GR has advisory roles with AstraZeneca, Pfizer, and Bayer in the past 2 years that have ended.

## Author contributions

BP – investigation, writing (original draft and editing); VM – methodology, formal analysis, writing (original draft and editing), RK – methodology, investigation, writing (original draft and editing), visualization; NDJ – methodology, investigation, writing (original draft and editing), visualization, data curation; SE – methodology, writing (editing), resources; DS – methodology, software, resources, writing (editing); RH – methodology, investigation, writing (editing); DEB - conceptualization, methodology, investigation, software, resources, data curation, writing (original draft and editing), visualization, supervision, project administration, funding acquisition.

All other authors are members of the ORIEN Microbiome Research Interest Group, with at least one member from each institution that contributed colorectal cancer cases to the study. BS, CHFC, MLC, RJR, SY, MRR, AAT, DPM, AM, TJB, RWL, HH, MNI, SH, CMU, JLR, GR, CDS – resources, writing (editing).

## Funding statement

This work was supported by the Holden Comprehensive Cancer Center ACS-IRG from the American Cancer Society (DEB). DEB was supported by the NIH, K08AI159619. RK was supported by the American Cancer Society (IRG-21-141-46-IRG) and the National Cancer Institute (R25CA273964). DS is supported by the National Institute on Aging (K01AG070310), the American Lung Association (1046611), and the American Cancer Society (RSG-23-1023205).

## Acknowledgements

We gratefully acknowledge the University of Iowa Holden Comprehensive Cancer Center’s Microbiome Core for their essential support in providing microbiome sequencing services. The Histology Research Laboratory at Iowa provided histology services.

---

**Table S1.**
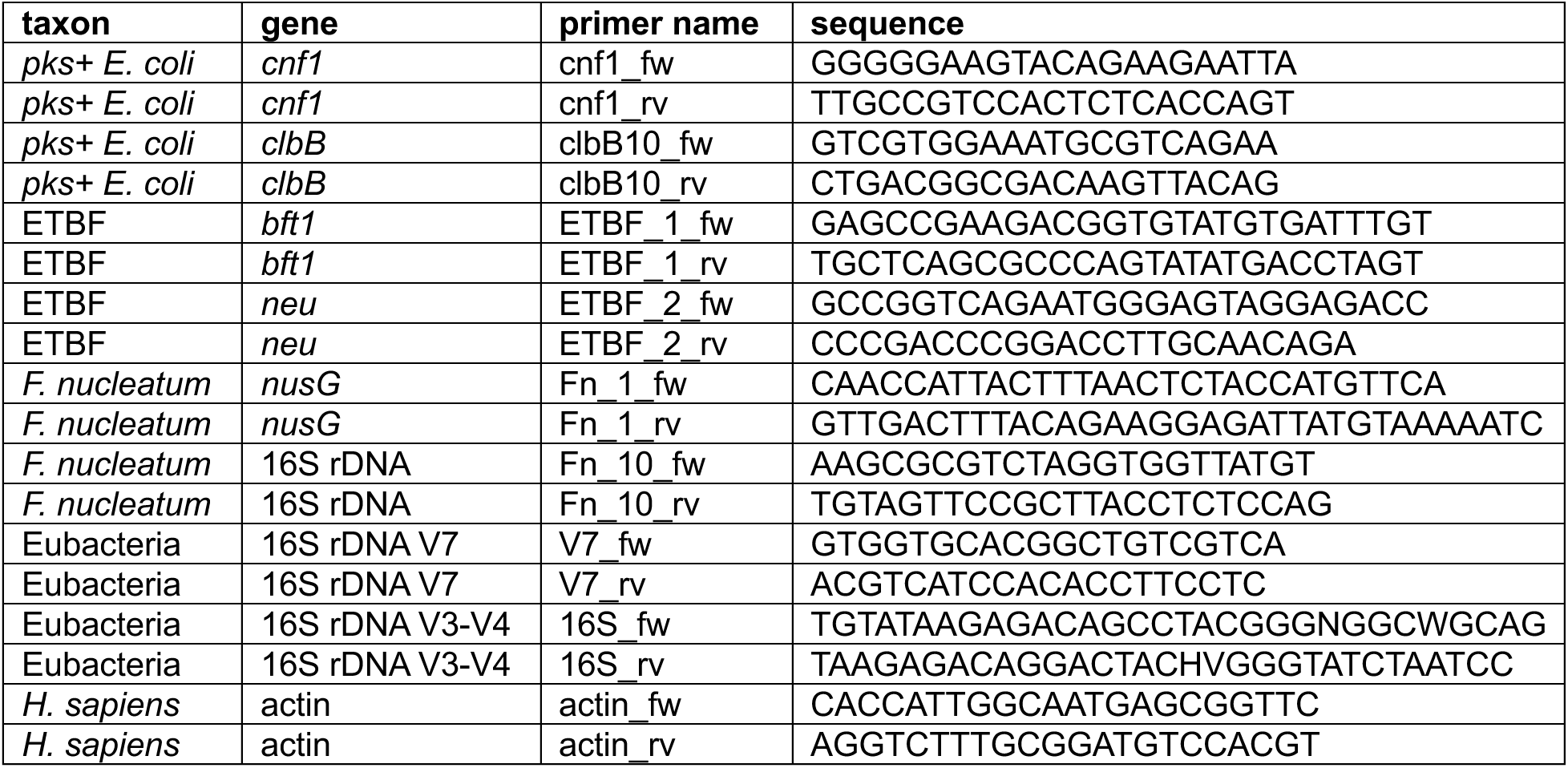
Quantitative PCR primers and targets for pathobiont-specific qPCR.

